# Rho activation drives luminal collapse and eversion in epithelial acini

**DOI:** 10.1101/2022.07.06.499064

**Authors:** Vani Narayanan, Purboja Purkayastha, Bo Yu, Kavya Pendyala, Sasanka Chukkapalli, Jolene I Cabe, Richard B. Dickinson, Daniel E. Conway, Tanmay P Lele

## Abstract

Epithelial cells lining a gland and cells grown in a soft extracellular matrix polarize with apical proteins exposed to the lumen and basal proteins in contact with the extracellular matrix. Alterations to polarity, including an apical-out polarity, occur in human cancers. While some aberrant polarity states may result from altered protein trafficking, recent observations of an extraordinary tissue-level inside-out unfolding suggest an alternate pathway for altered polarity. Because mechanical alterations are common in human cancer including an upregulation of RhoA mediated actomyosin tension in acinar epithelia, we explored whether perturbing mechanical homeostasis could cause apical-out eversion. Acinar eversion was robustly induced by direct activation of RhoA in normal and tumor epithelial acini, or indirect activation of RhoA through blockage of β1 integrins, disruption of the LINC complex, oncogenic Ras activation, or Rac1 inhibition. Furthermore, laser ablation of a portion of the untreated acinus was sufficient to induce eversion. Analyses of acini revealed high curvature and low phosphorylated myosin in the apical cell surfaces relative to the basal surfaces. A vertex-based mathematical model which balances tension at cell-cell interfaces revealed a five-fold greater basal cell surface tension relative to the apical cell surface tension. The model suggests that the difference in surface energy between the apical and basal surfaces is the driving force for acinar eversion. Our findings raise the possibility that a loss of mechanical homeostasis may cause apical-out polarity states in human cancers.

## Introduction

Most epithelial cells establish and maintain polarity along an apical-basal axis, that is, differences in the localization of membrane surface proteins at the apical and basal surfaces of the cell[1; 2]. Correct localization of apical and basal proteins is important for the physiological function of ductal, glandular, and tubular epithelial structures within organs[3]. Alterations to apical-basal polarity occur in several glandular cancers [4; 5; 6; 7; 8; 9]. For example, tumor epithelial spheres with “inverted” or “apical-out” polarity have been recently shown to collectively metastasize[10]. Yet, the mechanism by which tumor cells aggregates develop with apical-out polarity is not fully understood.

Epithelial cells derived from glandular tissues assemble spherical structures in 3D culture which are variously referred to as organoids, spheroids, enteroids or acini. These 3D acini feature a single hollow liquid-filled, pressurized lumen that are surrounded by the apical surface of the acinus, while the basal surface is adherent to the basement membrane assembled from extracellular matrix proteins [4]. Apical-basal polarity is a characteristic of epithelial spheroids, in which proteins like podocalyxin and ezrin localize exclusively to the apical surface[11; 12], while proteins like β1-integrins and β-catenin localize to the basolateral surfaces[13; 14; 15]. Protein sorting signals direct trafficking of apical and basal membrane protein containing transport vesicles leaving the trans-Golgi network to the respective membrane compartments[16]. Correct protein trafficking and membrane segregation of apical and basal proteins has been shown to be the principal mechanism for establishment of basal-out polarity during development of acini[17]. Additionally, the formation of tight junctions is required for restricting lateral diffusion of membrane proteins between the apical and basolateral membrane compartments[17; 18; 19].

Interestingly, a recent study by Co et al reported eversion from basal-out to apical-out polarity in acini formed from intestinal stem cells as they were transferred to suspension [20; 21]. The eversion was not caused by changes in protein trafficking but rather due to an extraordinary morphological rearrangement involving eversion of the apical surface and internalization of the basolateral surfaces. These observations raise the possibility that a similar tissue-level unfolding of acinar epithelia may cause apical-out polarity in cancers. RhoA-mediated actomyosin forces are an important contributor to epithelial acinar destabilization in cancer, and there is a positive correlation between increased actomyosin contractility and malignancy[22; 23]. Indeed, there are some similarities between the reported phenomenon of tissue eversion and previously hypothesized mechanical mechanisms for how contractility may promote outward migration of cells from within an acinar structure in the context of cancer[24]. In line with this, Co et al speculated that perturbations to mechanical forces may contribute to acinar eversion in their stem cell model [20]. Whether Rho mediated mechanical destabilization of normal or tumor epithelial acini can cause acinar eversion is unknown.

Cells in epithelial monolayers and in acini are polyhedral in shape with tetrahedral cell-cell lateral interfaces and polygonal apical and basal surfaces [25; 26]. Cell shape changes are driven by mechanical forces originating in the actomyosin cortex, located as a contractile sheet on cell boundaries. Surface tension from active contractile forces is opposed by cell adhesion (primarily mediated by E-cadherin), intracellular hydrostatic/osmotic pressure, as well as the opposing contractile force of neighboring cells. Cell shape in epithelial monolayers is governed by the relative surface tensions of the different interfaces [27; 28]. For example, higher tension in the lateral surfaces leads to flatter cells and lower tension leads to more columnar cells. Further, acinar homeostasis requires a mechanical balance of these forces to form a stable equilibrium (or quasi-static) state. As each cell in a typical epithelial acinus is contractile, there is a tendency for the acinus to reduce its area. A removal of the extracellular matrix can eliminate part of the resistance for a cell to contract, and this may contribute to the collapse of the acinar lumen, as has been observed when pre-formed acini are put into suspension. Further, such a collapse may create conditions favorable for eversion. This possibility is supported at least partially by the observation that blockage of β1-integrins, which mediate acinar adhesion to the basement membrane, can evert the apical surface while still in 3D matrix culture [21]. Likewise, we have previously reported that disrupting the LINC complex causes acinar collapse, and F-actin, which is typically at the apical surface, becomes present on the basal surface [29]. Therefore, here we explored whether perturbing mechanical homeostasis of acini can cause acinar eversion. Based on our findings, we propose a simple computational model which shows that a difference in apical and basal surface energy is the driving force for acinar eversion. Our results raise the possibility that eversion may be a potential mechanism for the generation of tumor spheroids with apical-out polarity.

## Results

### Rho activation drives E-cadherin dependent eversion of apical surfaces

We hypothesized that increasing cellular contraction in pre-formed acini should cause luminal collapse and lead to subsequent eversion of the apical surface. To test this hypothesis, we activated RhoA in pre-formed Madin-Darby canine kidney (MDCK) acini by treatment with Rho activator II that stabilizes RhoA in its GTP-bound form. The Rho effector, Rho kinase (ROCK), is then predicted to upregulate myosin light chain phosphorylation and increase actomyosin contractility, as we have recently shown in this model system [29]. Consistent with our hypothesis, treatment of control acini with Rho activator II caused a local breach in the acinar surface, followed by luminal collapse and dynamic eversion of the apical surface (Fig 1A, see Movies 1 to 3). Live imaging of acini containing cells expressing Emerald-occludin, which is present in tight junctions, and histone B-GFP, which localizes to nuclei, confirmed that cells moved *en masse* during the eversion while maintaining occludin localization in between cells in the everting front (Fig 1C, Movie 5). These features are similar to the apical-out eversion mechanism reported previously in suspended acini[20]. Such eversion was not observed in vehicle control treated acini (Figs 1B and 1E). Lumens of 25 out of 30 acini imaged with live cell imaging collapsed at a 1 μg/ml dose and underwent complete or partial apical-out eversion in the time-period of imaging. Consistent with Co et al[20], Rho activation altered the spatial distribution of apical podocalyxin not through effects on podocalyxin internalization and trafficking as in early acinar development[17] but rather through collective cell movement via breaches in the acinar walls while preserving the spatial segregation of podocalyxin at the cellular level.

**Figure 1.**
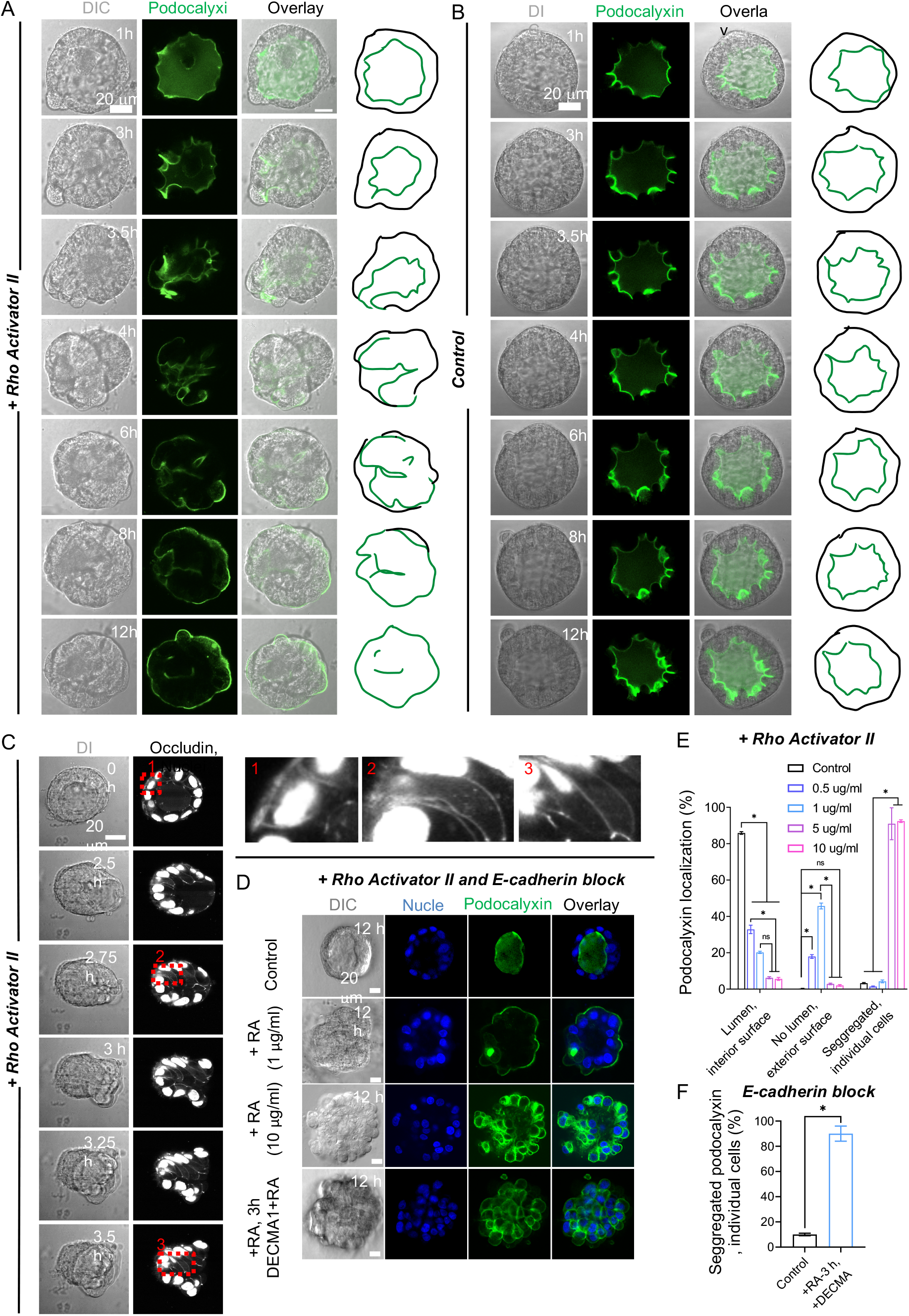
RhoA activation causes acinar eversion. Time-lapse differential interference contrast (DIC) images and confocal fluorescent GFP-podocalyxin images (GFP) of a 10-day old acinus treated with Rho activator (A) or treated with vehicle control (B) (see Movie 4). Scale bar, 20 μm. 1 μg/ml Rho Activator II was added at T = 0 h in A (see Movies 1 to 3). Images in A are representative of 25 acini out of the 30 assayed acini from three independent experiments (lumens of 5 acini did not collapse upon Rho activator treatment). Corresponding outlines show podocalyxin-labeled surface (green) and acinar outer contour (black). (C) Time-lapse DIC images and confocal fluorescent Emerald-occludin and GFP-H2B images (white) of a single acinus treated with Rho activator along with insets. 1 μg/ml Rho Activator II was added at T = 0 h (see Movie 5). Scale bar, 20 μm. (D) Confocal fluorescent images of a 10-day old control acinus, 10-day old acini treated with Rho Activator II at different doses for 12 h, and 10-day old acini treated with 1 μg/ml Rho Activator II (see movie 6) at 0 h and 10 μg/ml DECMA 1 at 3 h. Nuclei were stained with Hoechst 33342 (blue) and GFP-podocalyxin is shown in green. Scale bars, 20 μm. (E) Comparison of acinar frequency corresponding to the different Rho Activator II doses in D. Each data point is representative of 91 to 185 acini from 3 independent experiments. Error bars represent SEM, *p<0.05 as per ANOVA with Tukey’s multiple comparison test. (F) Comparison of disintegrated acinar frequency (with segregated podocalyxin around individual cells) corresponding to the DECMA 1 treatment in D. Each data point is representative of 100 acini from 3 independent experiments. Error bars represent SEM, *p<0.05 as per Student’s T test.

We found that the effect of RhoA activation on acinar eversion was dose dependent (Fig 1D). At higher doses of 5 μg/ml or 10 μg/ml of Rho activator II, eversion was less frequent. Instead, podocalyxin tended to become present over most or the entire surface of individual cells, precluding the possibility of collective podocalyxin surface movement and eversion (Figs 1D and 1E). The presence of podocalyxin over the entire cellular surface suggests a breakdown of the diffusive barrier to podocalyxin, which is maintained by tight junctions, likely due to a loss of mechanical coupling between cells. As E-cadherin-linkages transmit actomyosin contractile stresses between neighboring cells, and as treatment with Rho activator II upregulates actomyosin contractility in MDCK acini [29], we hypothesized that E-cadherin linkages between cells are unable to sustain the increased actomyosin contractile stresses at high doses of Rho activator II. Disruption of E-cadherin linkages would contribute to a general loss of cell-cell contacts and a loss of diffusive barriers to podocalyxin. We first induced luminal collapse by treatment with Rho activator II at the eversion-inducing dose (1 μg/ml), and then treated cells with DECMA-1, an E-cadherin function-blocking antibody, at the time (~3 h) when the apical surface should start to collectively move out of the acinus. Treatment of acini with DECMA-1 after 3 h of Rho activator treatment appeared to cause cellular separation, with podocalyxin again decorating the peripheries of individual cells, and preventing apical-out eversion (Fig 1D; bottom panel and Fig 1F). These findings are consistent with eversion occurring through collective cellular movement that requires E-cadherin cell-cell adhesions. RhoA activation at higher doses of Rho activator II caused luminal collapse but also likely disrupted E-cadherin cell-cell adhesions and tight junctions, preventing collective migration and causing loss of the diffusive barriers to podocalyxin between the segregated domains of the cell.

### LINC complex disruption causes inside-out acinar eversion

Our results above reveal that RhoA activation in a pre-assembled acinus is sufficient to cause apical-out eversion. To assess the robustness of these findings, we asked whether alternate methods to mechanically destabilize the acinus could also cause eversion. We previously reported that inducible disruption of the LINC complex in pre-formed MDCK acini causes luminal collapse and a shift in the localization of F-actin from the luminal surface to the external surface of acini reminiscent of the acinar eversion phenotype[29]. Here, we examined the extent to which LINC complex disruption altered the localization of podocalyxin. Podocalyxin was observed to localize on the entire exterior surface of the solid acinus after 72 hours of treatment with doxycycline to induce mCherry-KASH1 expression with a near complete loss in the central portion of the acinus (Fig 2A, bottom panel). The presence of podocalyxin on the entire exterior surface was observed for acini that lacked significant lumens (Fig 2A and 2B), and less commonly in acini that contained single or multiple lumens, consistent with our previous observations of exterior localization of F-actin in LINC-disrupted acini that lack lumens[29].

**Figure 2.**
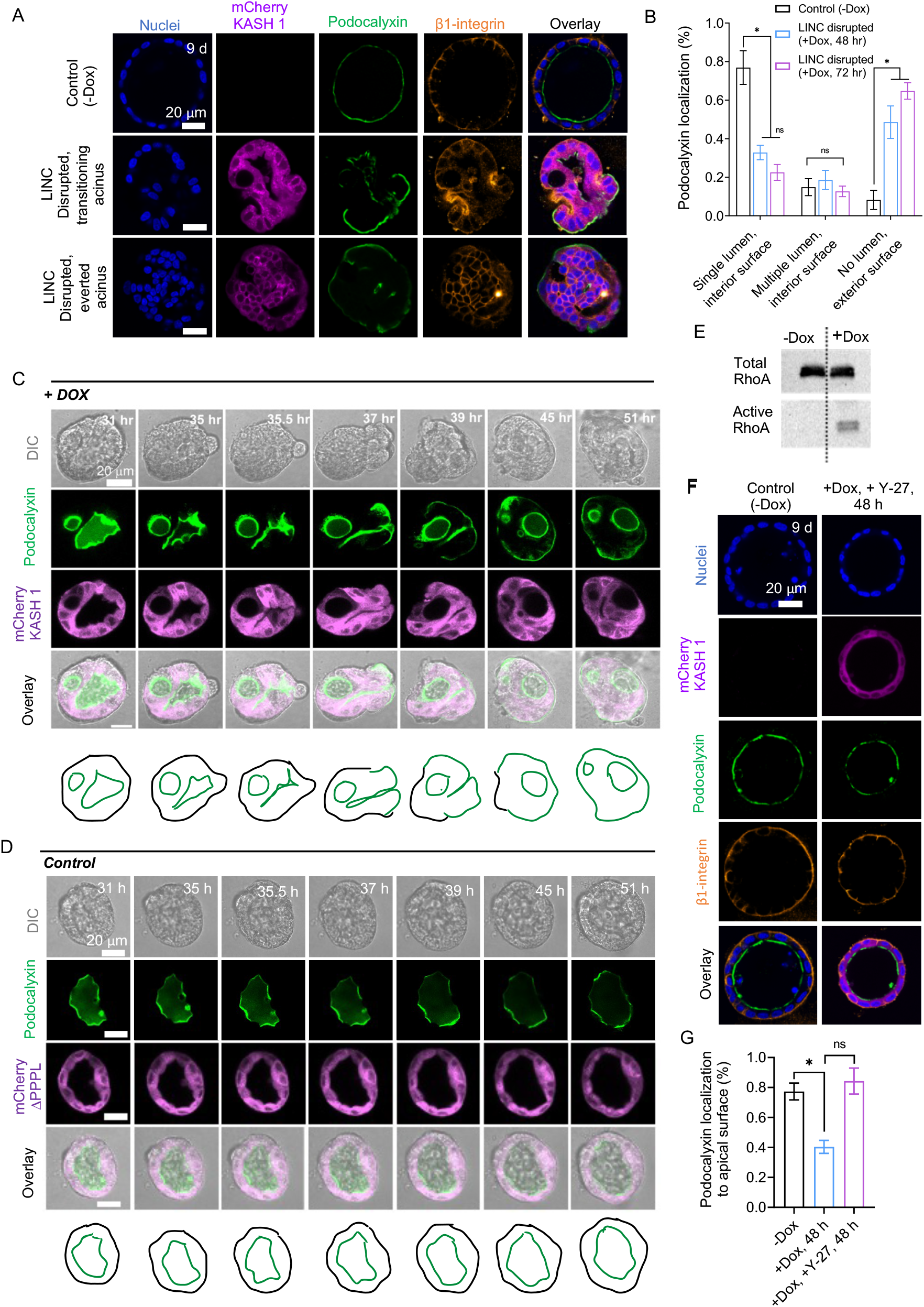
Indirect methods of Rho activation cause acinar eversion. (A) Confocal immunofluorescent images of 9-day old acini untreated with doxycycline (-Dox) or a 7-day old acinus treated with doxycycline for the indicated time (+Dox) to induce expression of mCherry-KASH1. Middle panel shows an acinus in the process of everting, while bottom panel shows a fully everted acinus. Nuclei were stained with blue. Scale bars, 20 μm. (B) Bar graph of the percentage occurrence of podocalyxin localization in untreated (-Dox) or treated (+Dox, 48 h, and +Dox, 72 h) acini, separately scored based on the presence of podocalyxin on the interior surface in acini with a single lumen, podocalyxin on the interior surface in acini multiple lumens, and podocalyxin on the exterior surface in acini with no lumens. At least 70 acini from three independent experiments were scored for each condition. Error bars represent ± SEM (*p < 0.05; One-way ANOVA test with post hoc Tukey [HSD] test). (C) Confocal fluorescent images of a 10-day old GFP-podocalyxin expressing acinus treated with 2 μg /ml doxycycline to induce mCherry-KASH1 expression at T = 0 h (see Movies 7 to 9), or (D) confocal fluorescent images of a 10-day old GFP-podocalyxin expressing acinus treated with 2 μg/ml doxycycline to induce mCherry-KASH1ΔPPPL expression (which does not disrupt the LINC complex); images are shown at the indicated times after this treatment (see Movie 10). The results in (C) represent 8 acini out of the 20 assayed acini from a total of three independent experiments (lumens in the 12 acini did not collapse over the time frame of the experiment). (E) Western blot shows total and active RhoA in absence and presence of doxycycline to induce mCherry KASH1 expression in MDCK cells in culture. The cells in 2D culture were treated with 100 μg/ml of doxycycline until a monolayer was formed, followed by the RhoA pulldown assay. (F) Confocal fluorescent images of GFP-podocalyxin and stained for β1-integrin of a 9-day old untreated acinus and a 7-d acinus treated with 40 μM Y-27632 for 48 h. DNA was stained with Hoechst 33342 (blue). Scale bar, 20 μm. (G) Comparison of acinar frequency with podocalyxin in the apical surface corresponding to the different conditions in (F). At least 70 acini from three independent experiments were scored for each condition. Error bars represent ± SEM (*p < 0.05; one-way ANOVA test with post hoc Tukey [HSD] test).

To test if the altered localization of GFP-podocalyxin was indeed due to apical eversion, we performed live imaging of GFP-podocalyxin localization in single MDCK-II acini. Acini were allowed to form for 10 days, and then treated with doxycycline to induce the expression of mCherry-KASH1. After 30 hours, we transferred the acini to a microscope and performed live imaging. We chose pre-formed acini in which GFP-podocalyxin was expressed clearly on the apical surface and seldom on the basal surface for imaging and imaged individual acini for up to 48 hours. Like the effects of Rho activation, mCherry-KASH1 expression resulted in eversion of the apical surface (Fig 2C, see Movies 7 to 9). In contrast, control acini assembled from cells engineered to express mCherry-KASH1ΔPPPL, which does not disrupt the LINC complex[29], did not collapse upon treatment with doxycycline (Fig 2D and Movie 10). Predictably, GFP-podocalyxin localization remained at the luminal surface upon expression of the mCherry-KASH1ΔPPPL construct.

Furthermore, we found that Rho was activated in LINC disrupted cells (Fig 2E). Inhibition of Rho kinase with Y27632 prevented acinar collapse in LINC disrupted cells as we have previously reported[29], as well as eversion (Fig. 2F and G). Blockage of β1 integrins or inhibition of Rac1 activation, both of which are known to also upregulate Rho activation in MDCK acini, also resulted in apical-out eversion phenotypes (Figs S1A, S1B and Movie 11). Similarly, doxycycline-induced expression of oncogenic H-Ras V12 which can also activate Rho[30] in pre-assembled acini also resulted in apical-out phenotypes (Fig S1C). These findings collectively show that apical eversion is a robust phenomenon that can be induced by multiple indirect approaches that upregulate Rho activity.

### Rho activation in tumor epithelial acini causes eversion

To explore whether a loss of mechanical homeostasis caused by Rho activation can similarly cause eversion in tumor epithelial acini, we examined the assembly of lung cancer cell tumor acini. We chose the metastatic Kras and p53 mutant mouse lung adenocarcinoma line 344SQ [31]. 344SQ cells cultured in 3D matrigel assembled acini with hollow lumens (Figure 3A). Treatment of 344SQ organoids with Rho activator II caused 344SQ acini to evert (Figure 3B and Movie 12), through a similar inside-out unfolding as observed for MDCK cells. Thus, Rho-driven eversion can occur in 3D structures assembled by normal and cancer cells, across different cell lines from distinct organs, and from different species.

**Figure 3.**
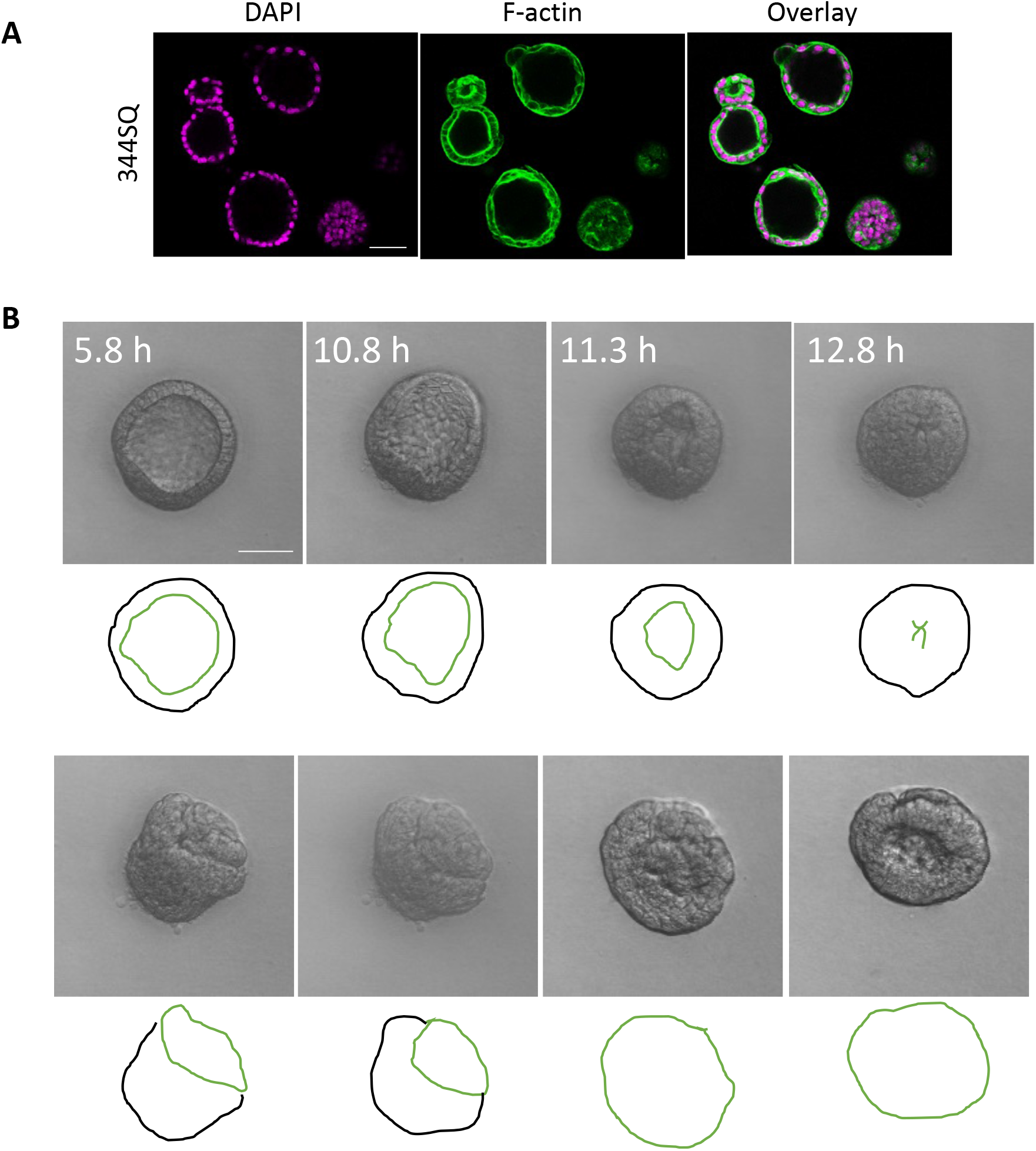
Rho activation causes eversion in tumor epithelial acini. A. Images of acini assembled by 344SQ tumor epithelial cells assembled in 3D Matrigel culture stained for F-actin or dNa. B. Time lapse images of a single acinus assembled by 344SQ cells, treated with Rho activator II (1 μg/ml) at 0 h (see Movie 12). Scale bar is 50 microns.

### Laser ablation of a portion of the acinar surface is sufficient to cause eversion

We interpret the effects of RhoA activation on epithelial acini as follows. Because RhoA activation increases Rho kinase mediated upregulation of actomyosin contractility, cells in the acini collectively try to reduce their cross-section. The resulting acinar contraction occurs through local breaches in the acinar walls which allows water to flow out of the lumen. To test this interpretation more directly, we ablated a portion of GFP-podocalyxin expressing MDCK acini cultured in 3D matrigel using femtosecond laser ablation (Fig 4A). Laser pulses from an 800 nm laser at a repetition rate of 80 MHz were focused on a spot size of roughly 20 μm. Laser ablation of the acinar wall was followed by a rapid squeezing of the luminal fluid out of the acinus and a collapse of the lumen (Fig 4B, Movie 13). The expulsion of the fluid from the lumen post ablation is consistent with the concept that the acinus is in a mechanical homeostasis with the tension in the acinar surface balancing the luminal pressure [32]. Importantly, ablation of a portion of the acinus resulted in an eventual apical eversion in a period of about 3 hours after the ablation (Fig 4C, Movie 14; six out of eight that were laser-ablated in this way underwent eversion; two out of eight rapidly disintegrated likely because the size of the spot was too large relative to the size of the acinus). These results show that luminal collapse can be triggered by directly disturbing the mechanical homeostasis of the acinus, and that this mechanical destabilization is sufficient to cause apical eversion.

**Figure 4.**
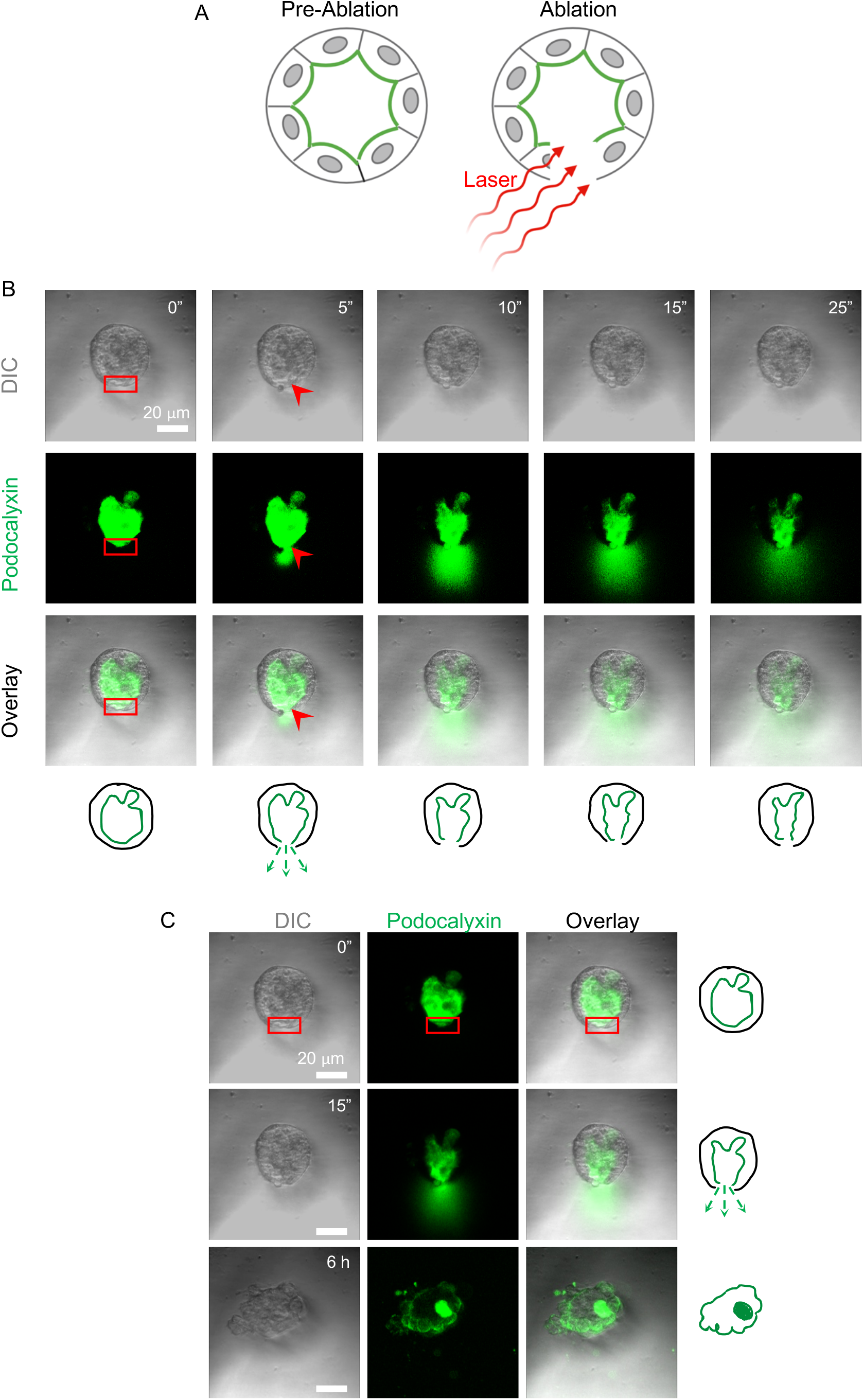
Laser ablation to disrupt mechanical homeostasis of acini. (A) Schematic of the laser ablation experiment; green color depicts podocalyxin. (B) Time lapse DIC and fluorescent GFP-podocalyxin images (GFP) of a 7-day old acinus that was ablated with the laser in the red box (see Movie 13); red arrow depicts damaged area from which fluid flows out. Corresponding outlines show podocalyxin-labeled surface (green) and acinar outer contour (black). Scale bar, 20 μm. (C) Time lapse fluorescent GFP-podocalyxin images (GFP) of a 7-day old control acinus post-ablation (see Movie 14). Scale bar, 20 μm. Corresponding outlines show podocalyxin-labeled surface (green) and acinar outer contour (black). Images in (C) are representative of 6 out of 8 acini.

### Apical surface tension is significantly lower than basal surface tension

In our experiments, we noted that the apical cell surfaces in the acini were concave facing the acinus center, despite the opposing lumen pressure, and their curvatures were consistently larger than the basal cell surfaces. We assume that such cell shapes in the acinar monolayer reflect a quasistatic force balance between neighboring cells, while also accounting for the pressure differences across the apical and basal cell interfaces. Absent any source or sink for radial flow across the monolayer, the intracellular hydrostatic pressure can be assumed uniform within a cell, and thus equal on apical and basal cell surfaces. Thus, a greater curvature of the apical surface suggests that its surface tension is less than that of the basal surface, according to the Law of Laplace. To quantify the differences, we measured the curvatures of the apical and basal surfaces of each cell from DIC or fluorescent images of Emerald-occludin (for basal surface radii) and fluorescent images of GFP-podocalyxin (for apical surface radii) of control acini as well as acini treated with Rho activator at the point of initiation of luminal collapse. We fit circles to each of the cellular surfaces to estimate the basal and apical surface radii of curvature, *r_o_* and *r_i_*, respectively (Fig 5A). We also fit circles to the overall acinus and to its lumen and calculated the outer and inner radii *R_o_* and *R_i_* respectively (Fig 5A). We found that the deviation of apical cell-surface curvature from the acinar luminal curvature was much larger than the deviation of the basal cell-surface curvature relative to the external acinar surface curvature (Fig 5B). Similar curvature differences persisted even upon Rho activation at the point of initiation of luminal collapse (Fig 5B).

**Figure 5.**
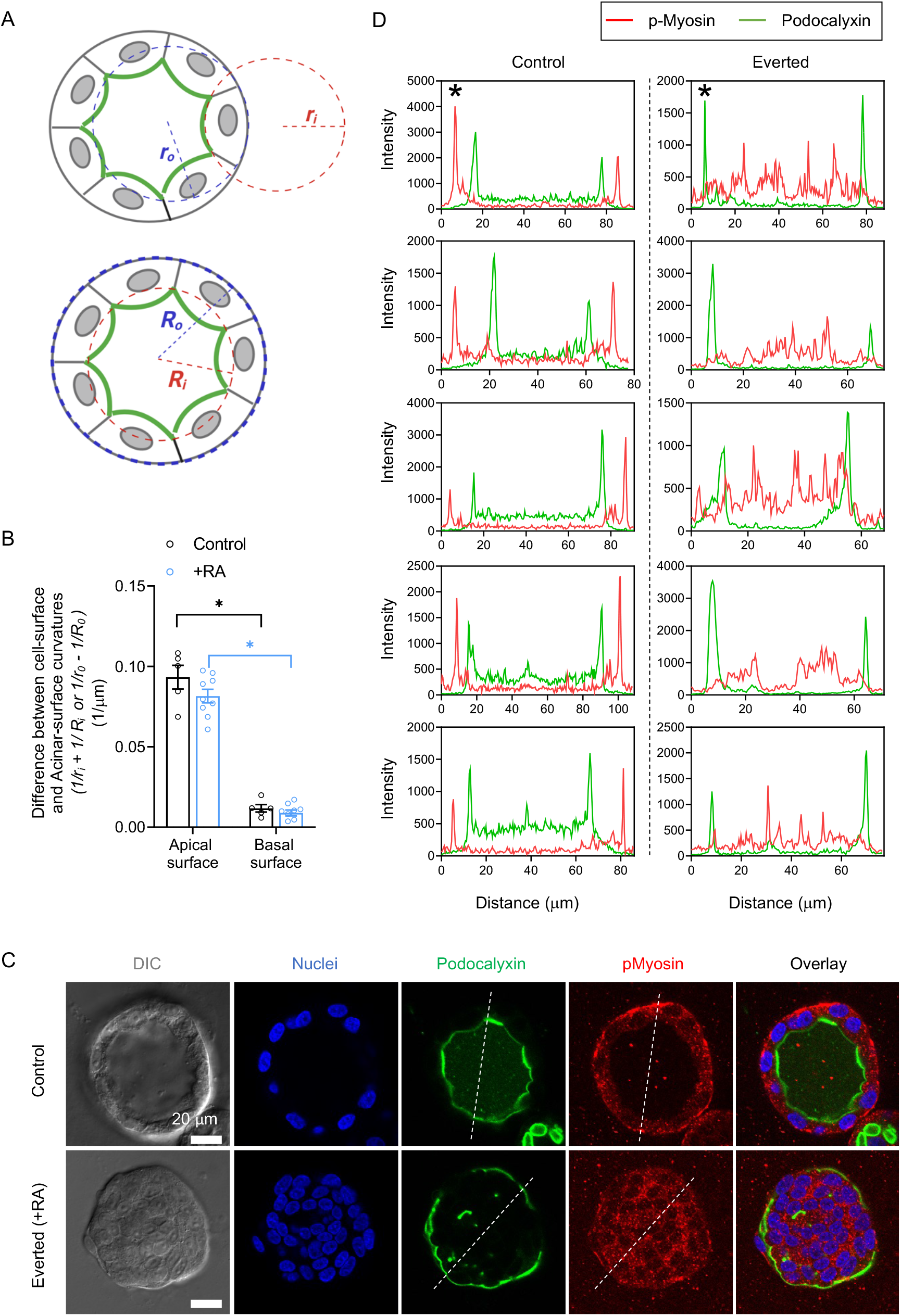
Acinar apical cell surface tension is lower than basal cell surface tension. (A) Schematics of circle fit method used for measuring radii of curvatures of apical cell surfaces (*r_i_)*, basal cell surfaces (*r_o_*), acinar inner radii (*R_i_)* and acinar outer radii (*R_o_)*. (B) Measured differences in curvature between the apical and basal cell surfaces (*1/r_i_* and *1/r_o_* respectively) and the acinar luminal and outer curvatures (*1/R_i_* and *1/R_o_*, respectively). The mean values are obtained from 5 control acini and 9 Rho activator treated acini with single lumen. Error bars represent ± SEM (*p < 0.05; as per Student’s T test). (C) Confocal fluorescent images of immunostained phosphorylated myosin and GFP-podocalyxin in control and an everted acinus; eversion was caused by treatment of a 10-day old control acinus with Rho Activator II at 1 μg/ml concentration. Nuclei are co-stained with Hoechst 33342 (blue). Scale bar, 20 μm. (D) Plots of intensities of podocalyxin (green curves) and phosphorylated myosin (red curves) along the diameter of each acinus (data from 5 control and 5 everted acini are shown). Asterisks mark the intensity plots along the indicated dashed white lines in the images in (C).

As phosphorylated myosin is a measure of cortical tension, we fixed control and everted (upon RhoA activation) acini expressing GFP-podocalyxin and stained them for phosphorylated myosin (Fig 5C). Phosphorylated myosin peak intensities occurred in the basal surface but not in the apical surface in control acini (Fig 5D; apical surfaces were detected by GFP-podocalyxin intensity peaks). Such peaks in intensity of phosphorylated myosin were absent in the exterior surface of everted acini and tended to be present inside the collapsed acinus. Collectively, these results further support our conclusion that the surface tension is lower in the apical surface compared to the basal surface, and that there is a tensional inversion in apical-out everted acini.

### Quantitative estimates of relative surface tensions in acinar surfaces

To quantitatively estimate the difference in the surface tensions between the apical surface and the basal surface from the observed curvatures, we formulated a mechanical model using a “mean-field” vertex model [33; 34] of a cell in the acinar monolayer mechanically connected with identical neighboring cells. As is commonly assumed in vertex models for cell monolayers, cell geometries are governed by the force balance between neighboring cells accounting for the cortical surface tensions of the cell interfaces. Elastic cell components are assumed to have relaxed on the relevant time scale or act in series with the cortical tensions without a primary impact on quasistatic cell shapes. In our model, we derive the force balance between a representative cell and its identical neighbors (Fig 6A) while accounting for the effect of surface curvature on the transmitted tension to neighboring cells. The representative three-dimensional cell is assumed to have a hexagonal cross section and curved apical and basal surfaces with radii of curvature *r_i_* and *r_o_*, respectively. As each cell in the acinar monolayer is connected to six identical neighboring cells, the surface tensions can act along apical (*τ_i_*), basal (*τ_o_*) and six lateral (*τ_c_*) surfaces of the cell. The inner (*R_i_*) and outer (*R_o_*) radii of the acinus are defined as the distance of the acinar center from corresponding inner and outer vertices along the lateral surfaces, respectively. *θ, η* and *φ* are angles defining the curved cell shape (see model details in methods section and Fig 6A).

**Figure 6.**
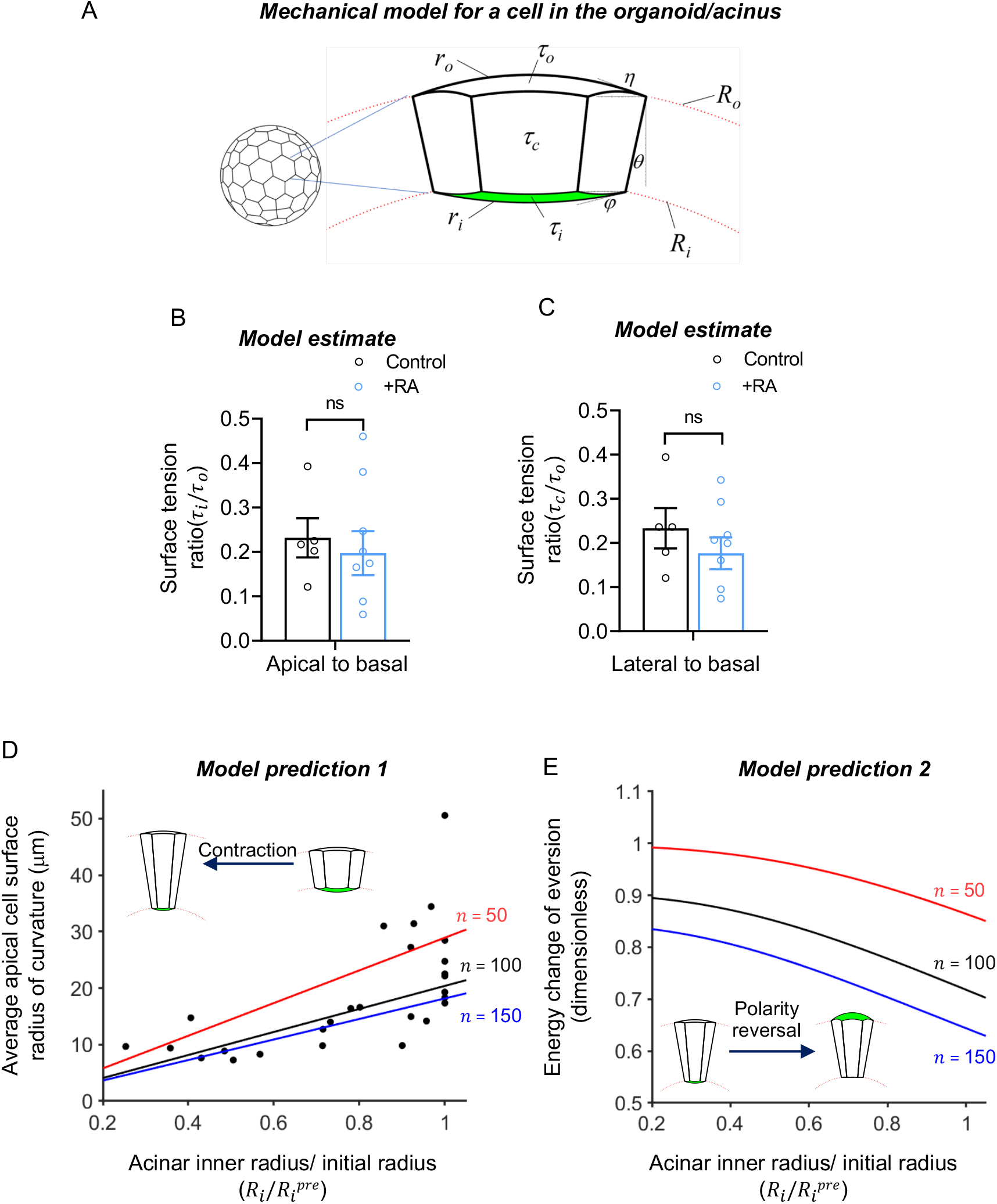
Model for acinar mechanics. (A) The model cell has a hexagonal horizontal cross-section and is interconnected with an identical neighboring cell. The acinus has inner and outer radii of curvature, *R_i_* and *R_o_*, respectively, defined by radial positions of the vertices. The apical and basal cell-surface radii of curvature, *r_i_*, and *r_o_*, respectively, are determined by the force balance at the vertices accounting for the contact angles, *φ* and *h*, and the surface tensions (*τ_o_*, *τ_i_* and *τ_o_* for the basal cell surface, apical cell surface, and cell-cell interface, respectively). (B) and (C) Apical-to-basal (*τ_i_/τ_o_*) and lateral-to-basal (*τ_i_/τ_o_*) surface tension ratios obtained from the curvature measurements and the corresponding equations (equations 7 and 6 respectively). The mean values are obtained from 5 control acini and 9 Rho activator treated acini with single lumen. Error bars represent ± SEM (ns represents p > 0.05 by Student’s T test). (D) Model predictions (lines) and experimental measurements (circles) of the apical cell-surface curvature versus radius (scaled to the initial radius) during acinar contraction following Rho activation. The lines are model predictions for different acinus cell numbers *n*. (E) Model predictions of energy change of eversion (calculated as initial surface energy of cell minus surface energy of everted cell using equation 8), which quantifies the driving force for eversion, versus relative acinus inner radius (same color scheme as (D)). The energy is normalized by *τ_o_V*^2/3^ where *V* is the cell volume.

Accounting for these geometric details, a force balance at the vertices yields formulae for the apical and lateral tensions relative to the basal tensions (equations in Figs 6B and 6C and in the model details in methods section). The formulae show that these relative tensions depend purely on the radii and angles. We experimentally measured the following parameters for each acinus containing a single hollow lumen: i) The inner (*R_i_*) and outer (*R_o_*) radii of the acinus, ii) the mean radii of curvature *r_i_* and *r_o_* of the apical and basal cell surfaces respectively, averaged over 6-8 cells in each acinar cross-section, iii) the angle *θ* calculated from the number of cells *n* in each acinus, which was in turn estimated from a relationship between *n* and *R_o_* (Fig S2 and methods). Substituting these measured parameters into the equation 6 and 7 (see methods) allowed calculation of the values of relative surface tensions *τ_i_/τ_o_* and *τ_c_/τ_o_* (Figs 6B and 6C and Fig S3) yielding average values of *τ_i_/τ_o_* ~ 0.23±0.04 and *τ_c_/τ_o_* ~ 0.23±0.05. Thus, the basal surface tension *τ_o_* is about four to five-fold larger than that of the apical surface *τ_i_* and the lateral surfaces *τ_c_*. These ratios were found to be roughly similar upon treatment for 2-3 h with Rho activator II (*τ_i_/τ_o_* ~ 0.20±0.05 and *τ_c_/τ_o_* ~ 0.18±0.04), suggesting that all surface tensions increased proportionately by Rho activation (Figs 5B and 5C; the measurements in Rho activator II treated acini were made just before the onset of acinar contraction).

Our experiments together with the model suggest that RhoA activation does not alter relative surface tensions even after treatment for up to 2-3 h, where the acini reach the point of luminal collapse. The model predicts that, if the relative surface tensions remain constant after the onset of luminal collapse, the cell apical surface radius of curvature (*r_i_*) will be proportional to the inner acinus radius (*R_i_*) during contraction, due to the forces of the lateral faces becoming concentrated on a smaller surface area and thereby increasing the intracellular pressure (see scheme in Fig S3). Consistent with this prediction, the experimentally measured apical surface radius of curvature (*r_i_*) of the acinus remained approximately proportional to the acinus inner radius during acinar contraction (circles in Fig 6D). The lines show model predictions for the dependence of *r_i_* on acinus radius for relative surface tensions assumed constant at *τ_i_/τ_o_* ~ 0.23±0.04 and *τ_c_/τ_o_* ~ 0.23±0.05 (see Figs 6B and 6C) parameterized by the number of cells. The agreement between the model and the data validates the model assumptions regarding surface tension balancing a uniform intracellular pressure and suggests that the relative surface tensions remained constant during the initial phase of contraction.

### Difference between apical and basal surface energy increases during acinar contraction

Based on the observations that i) phospho-myosin intensity is lower in the acinar apical surface than the basal surface in control acini, ii) eversion results in an exchange of the phosphomyosin-labeled apical surface with smaller area with the phosphomyosin-labeled basal surface with larger area, and iii) apical surface curvatures are higher than basal surface curvatures, we hypothesized that a difference in the apical and basal surface energies is the driving force for eversion. We used the model to calculate the total surface energy in a control acinar cell from the numerically integrated surface areas of the various surfaces of the model cell multiplied by the corresponding constant relative surface tensions, *τ_i_/τ_o_* ~ 0.20±0.05 and *τ_c_/τ_o_* ~ 0.18±0.04 (see Equation 8). Likewise, the energy of the everted acinar cell at the same acinar inner radius (with the basal and apical surface tensions switched) was calculated, and the energy difference was interpreted as the driving force for eversion. The model predicts that the energy change or driving force for eversion increases as the acinar inner radius decreases during contraction of the acinus following Rho activation (Fig 6E). As the acinus contracts to smaller radii, the apical surface area decreases faster than the basal surface area (assuming constant cell volume). Consequently, the driving force for the apical-out polarity state becomes stronger with the decreasing acinus radius (Fig 6E). Thus, the model predicts an increasing driving force for apical-out polarity during luminal collapse induced by Rho activation. This increased driving force explains the tendency toward apical-out eversion at smaller luminal radii.

## Discussion

It has been shown recently that glandular cancer cell spheroids with apical-out polarity invade tissue and contribute to cancer metastasis[10]. The mechanism by which such spheroids develop apical-out polarity is unclear. An upregulation of RhoA-regulated actomyosin tension occurs in glandular cancers[35], but whether such changes can promote apical-out polarity is not known. Here we found that RhoA activation causes a collapse of the lumen and subsequent apical-out eversion in in epithelial acini, including acini assembled by tumor epithelial cells. Our measurements and theoretical model support a mechanism in which a difference in surface tension between the apical and basal surfaces favors acinar eversion. That is, eversion involves transitioning from a higher-energy metastable state to a lower-energy state of reversed cell polarity. Laser ablation of the acinus is sufficient to allow this transition, indicating that Rho activation is not a necessary condition for acinar eversion. Rather, Rho activation facilitates eversion by allowing the acinus to contract due to a breach in the walls of the acinus that allows the luminal volume to reduce through an efflux of the luminal water. Acinar contraction results in an even lower apical area compared to the basal area of the now elongated acinar cells, corresponding to an even lower surface energy in the apical surface compared to the basal surface of the cells (surface energy being the product of tension and area). In this process, the five-fold difference in surface tension in the basal to apical surface does not appear to change. The increased driving force upon acinar contraction caused by a reduction in the apical area relative to the basal area is what drives eversion.

Because of the lower energy state after eversion, it would be unlikely for acini to spontaneously undo eversion and revert through a similar outside-in cohesive movement of cells. Supporting this, acini everted by mCherry-KASH1 LINC complex disruption were unable to return to a normal basal-out phenotype when mCherry-KASH1 expression was turned off (data not shown). This may be important in the context of human diseases, such as cancer, as it may be unlikely for cells in an apical-out phenotype to return to correct basal-out polarity.

Our observation that apical-out polarity is established through eversion rather than changes in protein trafficking contrasts with several prior studies by Mostov and others [2; 3; 36; 37] showing that RhoA-induced apical-out polarity occurs because of altered trafficking of apical and basal proteins. An important difference is that our experiments involved perturbations to pre-formed, well-established hollow structures, whereas prior studies primarily studied the effect of RhoA perturbations during acinar development[38]. Blocking β-1 integrins and inhibition of Rac-1, two treatments previously shown to affect apical-basal protein trafficking during acinar development[17; 37], instead resulted in eversion when applied to pre-formed acini. Thus, there appear to be different mechanisms by which apical-basal polarity changes occur, which are dependent on the state of the acinus or organoid (developing vs well-formed).

Eversion of glandular acinar structures with hollow lumens may be a potential mechanism by which tumor spheres with apical-out polarity assemble *in vivo* [10], and subsequently collectively metastasize. Rho activation in acinar cells in response to a stiffening of the extracellular matrix [22; 35; 39; 40; 41] may cause a switch from the hollow luminal phenotype to an everted phenotype. Alternatively, tumor eversion could be driven by a downregulation of integrin binding or the expression of oncogenic H-Ras, both of which caused eversion in our experiments. Overall, our results suggest that apical-out polarity in epithelial acini is a robust phenomenon that may be a contributing factor in tumor progression.

## Materials and Methods

### Stable cell lines and cell culture

MDCK cells, a gift of Rob Tombes (VCU), were used in these studies. Generation of MDCK cell lines stably expressing doxycycline-inducible mCherry-DN-KASH1 or mCherry-KASH1ΔPPPL was previously described[42]. To generate cells expressing GFP-podocalyxin, a retrovirus was generated using pQCXIN GFP-podocalyxin (gift of Ira Mellman[34]), and cells were selected based on fluorescent expression. MDCK cells stably expressing occludin-emerald and H2B-eGFP were a gift of Teemu Ihalainen. To generate MDCK cells stably expressing a doxycycline-inducible GFP H-Ras V12, lentivirus was generated using pcDNA/TO/GFP H-Ras V12 (gift of Yasuyuki Fujita)[43] and pLenti CMV TetR Blast (Addgene plasmid 17492), and cells were selected based on fluorescent expression.

MDCK cells were cultured in DMEM medium with 4.5 g/l glucose (ThermoFisher), supplemented with 10% v/v fetal bovine serum (FBS, ThermoFisher) and 1% v/v penicillinstreptomycin mix (Pen-Strep, ThermoFisher). Lung cancer cells (344SQ stably transfected with a non-coding siRNA) were a kind gift of Dr. Jonathon M Kurie at mD Anderson Cancer Center. The lung cancer cells were cultured in RPMI medium (ThermoFisher), supplemented with 10% v/v fetal bovine serum (FBS, VWR) and 1% v/v penicillin-streptomycin mix (Pen-Strep, ThermoFisher).

For 3D acinar culture, cells were trypsinized from tissue culture plates and 5000 cells were suspended in 400 μl growth medium supplemented with 2% v/v Matrigel. The cell suspension was seeded onto an 8-well Nunc Lab-Tek II chambered coverglass or coverslip; pre-coated with 40 μl of growth factor reduced (GFR) Matrigel (Corning) as previously described[42]. The pre-coated Matrigel was allowed to solidify by incubation for at least 30 min at 37°C before seeding the cells. The growth medium was changed every 3 to 5 days and the cells were allowed to form acini for 7-12 days [4].

### LINC complex disruption and Drug Treatment

Acini were treated with 1 and 2 μg/ml doxycycline (EMD Millipore) for up to 72 hours to induce dominant negative mCherry-KASH1 expression for LINC complex disruption. In other experiments, acini were treated with Rho activator II (Cytoskeleton Inc) at final concentrations of 1, 5 and 10 μg/ml for 12-20 h or Y-27632 (Cayman Chemical) at final concentration of 40 μM for 48 h.

### Blockage of β-1 Integrins, E-cadherins and inhibition of Rac-1

Acini were treated with anti-integrin β1, clone AIIB2 antibody (Millipore Sigma) at final concentration of 8 μg/ml for 12 h. Acini were treated with Anti-E cadherin antibody, clone DECMA-1 (Abcam) at final concentrations of 10 μg/ml for 9 h following Rho Activator II treatment at final concentration of 1 μg/ml for 3 h. Rac-1 was inhibited in acini by treating them with NSC-23766 (APExBIO) at final concentration of 50 μM for 24 h.

### Immunofluorescence staining

Immunostaining of acini was performed as previously described^8^. Briefly, acini were fixed with 2% paraformaldehyde for 1 hour at 37°C and rinsed thrice with PBS-Glycine (0.1M), with a 5 min interval between each rinse. Then, the acini were permeabilized with a permeabilization buffer (0.5% Triton X-100 in PBS) for 30 min at room temperature. This was followed by incubation with an IF buffer (130 mM NaCl; 7 mM Na2HPO4; 3.5 mM NaH2PO4; 7.7 mM NaN3; 0.1% BSA; 0.2% Triton X-100; 0.05% Tween-20) that was supplemented with 10% goat serum at room temperature. Next, the acini were incubated with the primary antibody mouse anti-podocalyxin (EMD Millipore, clone 3F2:D8, working dilution 1:200), mouse β1 integrin (Invitrogen, CD29, working dilution 1:250), rabbit monoclonal anti phospho-myosin light chain (Ser 19) (Cell signaling, working dilution 1:50) or rabbit monoclonal anti E-cadherin (24E10) (Cell Signaling, working dilution 1:1600) overnight at 4°C. The acini were then rinsed with PBS and incubated with secondary antibody Alexa Fluor 647 goat anti-mouse antibody (Invitrogen, working dilution 1:200), or Alexa Fluor 488 donkey anti-mouse antibody, or Alexa Fluor 594 goat anti-rabbit antibody (Invitrogen, working dilution 1:250) or Alexa Fluor 647 goat anti-rabbit antibody (Invitrogen, working dilution 1:250) for 1 hour in the dark at room temperature. Phalloidin conjugated to a fluorophore (Alexa Fluor 594 and Alexa Fluor 488, Life Technologies) was used to stain F-actin and Hoechst 33342 (Life Technologies/Invitrogen) to stain the nucleus. The immunostained acini were mounted on a glass coverslip using Vectashield Antifade mounting medium (Vectorlabs) with DAPI (when Hoechst 33342 is not used) or without DAPI before imaging.

### Confocal imaging

For some experiments, live cell confocal imaging of acini was performed under a Nikon Ti2 Eclipse A1+ confocal microscope (Nikon, USA) using Nikon CFI Apochromat LWD Lambda S 40X/1.15 NA water immersion objective including Nikon TI2-N-WID water immersion dispenser to prevent water evaporation. Other live cell imaging experiments were performed under Olympus FV3000 (Olympus Scientific Solutions Americas Corp., USA) using Super Apochromat 30X/1.05 NA silicone oil immersion objective (UPLSAPO30XSIR) or under Zeiss LSM 900 with Airyscan 2 (Carl Zeiss Microscopy, USA) using LD C-Apochromat 40X/1.1 NA W Corr M27 water immersion objective (421867-9970-000). The correction collar of the objective was adjusted based on the manufacturer’s suggestion for different cover glasses. In all live-cell imaging experiments, acini were maintained in environmental chamber (*In Vivo* Scientific or Tokia Hit or Zeiss respectively) at 37°C and 5% CO2. Fixed cell fluorescence imaging was performed under Olympus FV3000 (Olympus Scientific Solutions Americas Corp., USA) using Super Apochromat 60X/1.3 NA silicone oil immersion objective (UPLSAPO60XS2) or using Plan Apochromat 20X/0.8 NA air objective ; or under Zeiss LSM 900 with Airyscan 2 (Carl Zeiss Microscopy, USA) using W Plan Apochromat 20X/1.0 NA objective (421452-9681-000) or LD C-Apochromat 40X/1.1 NA W Corr M27 water immersion objective (421867-9970-000); or under a Zeiss LSM 710 laser scanning (Carl Zeiss Microscopy, USA) using Plan Apochromat 40X/1.1 NA water immersion objective. For all confocal imaging, a pinhole opening of 1 Airy disk was used.

### Laser ablation

Laser ablation of acini was performed by focusing energy from a Ti: Saphire (720-950 nm) pulsed laser (Coherent Chameleon Ultra) over a spot size of 20 μm through the objective lens at a wavelength of ~800 nm, nominal laser head power of 2.5W, pulse duration of 140 fs and a repetition rate of 80 MHz. Time lapse images over 12 hours were acquired under a Zeiss LSM 780 NLO multiphoton microscope (Carl Zeiss Microscopy, USA) using LD C-Apochromat 40X/1.1 NA W Corr M27 water immersion objective (421867-9970-000). The imaging was performed at 37°C and 5% CO2 using a temperature-controlled motorized stage.

### RhoA activation pull down assay and western blotting

The RhoA Activation Assay Biochem Kit (bead pull-down format, Cytoskeleton) was used for performing the pull-down assay on 2D monolayers that were either uninduced or induced for the (DN) KASH1 mutation. The experiment was performed with the help of the manufacturer’s protocol as described here forth. (DN) KASH1-inducible cells that were sub-cultured, induced for LINC disruption at the time of seeding with 100 μg/ml of doxycycline, and incubated at 37°C until monolayer confluency was reached, were compared against the non-induced control monolayers. Monolayers were washed with ice-cold phosphate-buffered saline (PBS) and kept on ice. Cells were lysed using the Cell Lysis Buffer (50mM Tris pH 7.5, 10mM MgCl2, 0.5M NaCl, and 2% Igepal) supplemented with 1% v/v protease inhibitor cocktail (62 μg/ml Leupeptin, 62 μg/ml Pepstatin A, 14 mg/ml Benzamidine and 12 mg/ml tosyl arginine methyl ester) on ice. Following the harvest of the lysates with a cell scraper, the supernatant was collected, snap-frozen, and preserved at -80°C. About 20μl of the lysate was isolated in order to measure protein concentrations with the help of Pierce™ 660nm Protein assay Reagent and Bio-Rad SmartSpec Plus Spectrophotometer.

For the assay, the lysates were thawed on ice, equalized to a protein concentration of ~1mg/ml, of which 50 μg was saved for Western Blot quantification of total RhoA, and 400 μg was used for the non-hydrolysable GTP analog (positive cellular protein control, GTPγS). Approximately 400 μg of each sample lysate, including GTPγS, which was prepared for loading as per the manufacturer’s protocol, were immediately incubated with rhotekin-RBD beads (50 μg) for 1h at 4°C on a rotator. After incubation with the beads, the samples were centrifuged at 5000 x g for 1 min at 4°C, following which 90% of the supernatant was carefully removed and the beads were washed once with the Wash Buffer (25 mM Tris pH 7.5, 30 mM MgCl2, 40 mM NaCl) and centrifuged at 5000 x g for 3 min at 4 °C. The pellet from the bead samples were thoroughly resuspended in 2X Laemmli Sample Buffer supplemented with 5% ß-mercaptoethanol, boiled for 2 min, and run on a 4-20% bis-acrylamide crosslinked gel along with 20ng of the His-RhoA control protein, 50 μg of the total lysate for assessing total RhoA, and the positive cellular protein control (GTPγS). After electro-blotting, the protein was transferred to a polyvinylidene fluoride (PVDF) microporous membrane for 90 min at 90 V and 4 °C. The membrane was then washed with Tris-buffered Saline (TBS 1X), dried, rehydrated, and blocked with a blocking buffer (5% BSA, 10 mM Tris-HCl pH 8.0, 150 mM NaCl, 0.05% Tween 20) for 1 hour at RT. The membrane was then probed with a mouse monoclonal anti-RhoA antibody (supplied with the Cytoskeleton Kit) at a working dilution of 1:500 overnight at 4 °C with constant agitation. After multiple membrane washes with TBST (10 mM Tris-HCl pH 8.0, 150 mM NaCl, 0.1% Tween 20) at RT, the membrane was incubated with an anti-mouse HRP-conjugated secondary antibody (Cell Signaling, working dilution of 1:5000) for 1 hour at RT with constant agitation. The RhoA signal was then observed using the BioRad ChemiDoc Touch Gel Imaging System. To assess for an additional loading control, the membrane was washed with TBST, stripped using a stripping buffer (Thermofisher), and probed with a mouse anti-tubulin antibody (clone dm1a, Sigma, working dilution of 1:5000) overnight at 4 °C with constant agitation. After multiple membrane washes with TBST at RT, the membrane was incubated with an anti-mouse HRP-conjugated secondary antibody (Cell Signaling, working dilution of 1:10000) for 1 hour at RT with constant agitation. The tubulin signal was then observed using the BioRad ChemiDoc Touch Gel Imaging System.

### Image analysis

The number of acini corresponding to various treatment conditions in Figs 1E, 1F, 2B, 2G and Figs S1A and S1B were counted using the multipoint tool in ImageJ. The phosphorylated myosin intensities in control and everted acini (Fig 5D) were obtained by measuring the radial intensity profile in ImageJ along a straight line drawn using the line tool in ImageJ across any diameter of the acinar cross-section (white dashed lines in Fig 5C). All the radii of curvatures in Figs 5B and 5D were obtained using “circle-fit” method (Fig 5A) where a circle was manually drawn using circle tool of ImageJ such that the circle fits the corresponding cell or acinar surface. The surface area of each circle was calculated in ImageJ and the corresponding radii of curvatures were obtained. Other measurements required for the model parameters like number of cells per acini (*n)*, acinar outer surface area in Fig S2 and acinar volumes were also calculated from confocal z-stacks of acini using several ImageJ tools or Plug-ins. The time lapse images were converted to movies using ImageJ.

### Statistical analysis

The statistical comparisons between control and treatment groups were done using the Analysis of Variance (One-way ANOVA) test followed by a further comparison of the groups using the Tukey (HSD) test (Fig 1E, 2B and 2G) and the Student’s T test assuming equal variances (Fig 1F, 5B, 6B, 6C, bar graphs in Figs S1A and S1B). All statistical tests were conducted at a 5% significance level (*p < 0.05). Detailed experimental design and statistical information can be found in Figure Legends. Graphical plots were generated using GraphPad Prism version 9.3.1 (GraphPad) and MATLAB version R2022a (Mathworks). Schematics were generated using BioRender and MATLAB version R2022a (Mathworks).

### Mathematical model

#### Cell Geometry

A representative cell in the acinus monolayer was modeled as a three-dimensional cell with a hexagonal cross-section and curved apical and basal surfaces with radii of curvature *r_i_* and *r_o_*, respectively. The model cell is connected to six identical neighboring cells in the curved monolayer (Fig 6A). The stresses that govern the cell shape at mechanical equilibrium are generated by surface tensions along the apical surface (*τ_o_*), the basal surface (*τ_i_*), and the six cell-cell interfaces (*τ_c_*). The acinus inner and outer radii, *R_i_* and *R_o_*, respectively, are defined at the vertices connecting the surfaces. Let *γ_o_* be the horizontal distance from a vertex at *R_o_* to the center of the hexagonal cross-section (of area 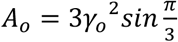) connecting the six vertices. The angle *θ* of cell edge relative to vertical direction is given by:

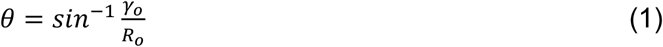

For an acinus consisting of *n* cells, the acinus outer surface area is approximately *nA_0_ = 4πR_o_^2^*. Thus, *θ* can be written as:

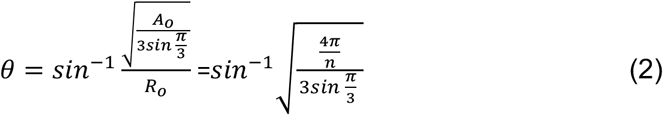

The parameters *R_i_*, *R_o_*, and *θ*, together with the basal and apical cell-surface curvatures, *r_o_* and *r_i_*, respectively, fully define the shape of the model cell. The parameters are constrained by the cell volume *V*, which is assumed constant. An average cellular volume of V ~1500 μm^3^ was obtained from the experimentally measured acinar volume, measured lumen volume and the measured cell numbers. The angle *θ* was found to depend on the number of cells *n* in the acinus (Equation 2). To calculate the number of cells in the acinus, we performed 3D confocal fluorescence microscopy of acini with co-stained nuclei and measured the number of cells (*n*) in each acinus by counting the total number of nuclei across the z-stacks. We found that *n* was directly proportional to the acinar outer surface area (*n* = 4*πkR_o_^2^*). We obtained the value of the proportionality constant (*k* = 4.07×10^-3^ μm^- 2^) from our measurements (Fig S2) and used this value to obtain values of *n* corresponding to measured values of *R_o_* for all control acini as well as acini treated with Rho activator II just before luminal collapse. The values of *n* were used to obtain corresponding values of *θ* (from Equation 2 in methods) for each acinus.

#### Cell Mechanics

The surface tensions of the basal cell surface (*τ_o_*), the apical cell surface (τ_i_), and cellcell interfaces (*τ_c_*) can be related to the above geometric parameters as follows. The angle between the hexagonal cross section at *R_o_* and cell surface at a vertex is

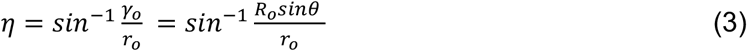

Similarly, the corresponding angle at *R_i_* is

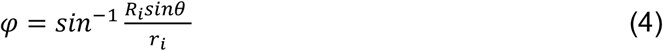

Accounting for the angles of interconnecting surfaces, the force balance at the vertices, yields

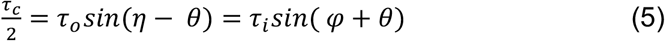

Combining Equations. 3-5 gives equations relating the surface tensions to the geometry:

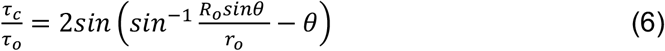

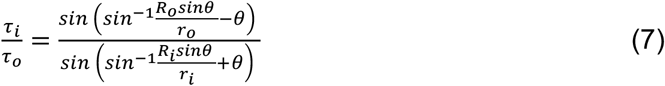

Note that for fixed values of the surface tensions, the ratios 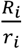 and 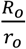 are constants for a given *θ* for any acinus radius. The total surface energy of the cell is given by

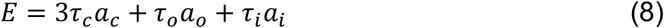

where *a_c_*, *a_o_*, and *a_i_* are the surface areas of the cell-cell interface, the (curved) basal and apical surfaces, respectively. The surface areas were calculated numerically for a given geometry.

## Acknowledgments

We thank Dr. Gregg Gundersen for his helpful suggestions. This work was supported by NIH U01 CA225566 (T.P.L. and R.B.D.) and a CPRIT established investigator award grant # RR200043 (T.P.L.). DEC acknowledges support from NIH R35GM119617 and NSF CAREER award CMMI 1653299.

## Competing Interests

The authors have NO competing interests

## Notes

### Competing Interest Statement

The authors have declared no competing interest.

### Summary of Updates

New data has been added and the manuscript revised substantially.

