## Supplemental figures and movie legends for "Rho activation drives luminal collapse and eversion in epithelial acini"

**Supplementary Figures**

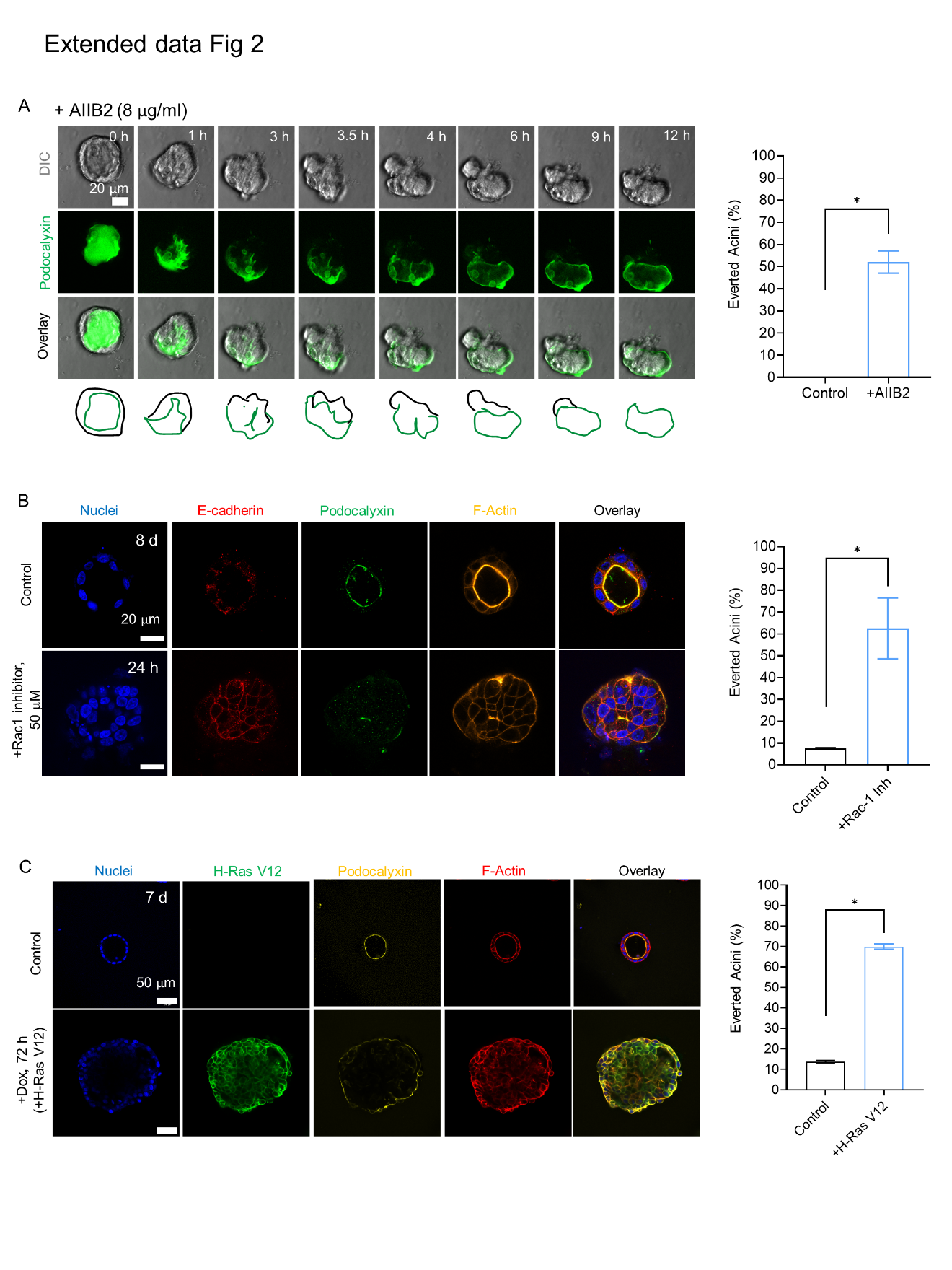

**Fig. S1.** (A) Time-lapse differential interference contrast (DIC) microscopy images and fluorescent GFP-podocalyxin images (GFP) of a 7-day old acinus treated with AIIB2 (β-1 integrin function blocking antibody). 8 μg/ml AIIB2 was added at T = 0 h (Movie 11). About 50% of inhibitor-treated acini everted over 12 hours. Data is representative of total of 65 control acini and 132 AIIB2 treated acini from 3 independent experiments. Error bars are SEM, *p<0.05 by Student’s T test. Scale bar: 20μm (B) Percentage of everted MDCK II acini with and without Rac1 inhibitor (Rac1 inh, 50μM) treatment on day 7 of acinar morphogenesis, for 24 hours. About 60% of inhibitor-treated acini everted over 24 hours. Data is representative of a total of 218 acini (untreated control) and 250 acini (Rac1 inh, 24h) over 2 independent experiments (*p < 0.05, Student t-test). Scale bar: 20μm (C) MDCK cells expressing doxycycline-inducible GFP H-Ras V12 were grown in Matrigel for 7 days. Doxycycline was added to the 7-day old acinus at T=0 to induce GFP H-Ras V12 expression, for 72 hours. About 70% of doxycycline induced acini everted after 72 hours. Data is representative of 60 acini per condition from 3 independent experiments. Error bars are SEM, *p<0.05 by Student’s T test. Scale bar: 50μm

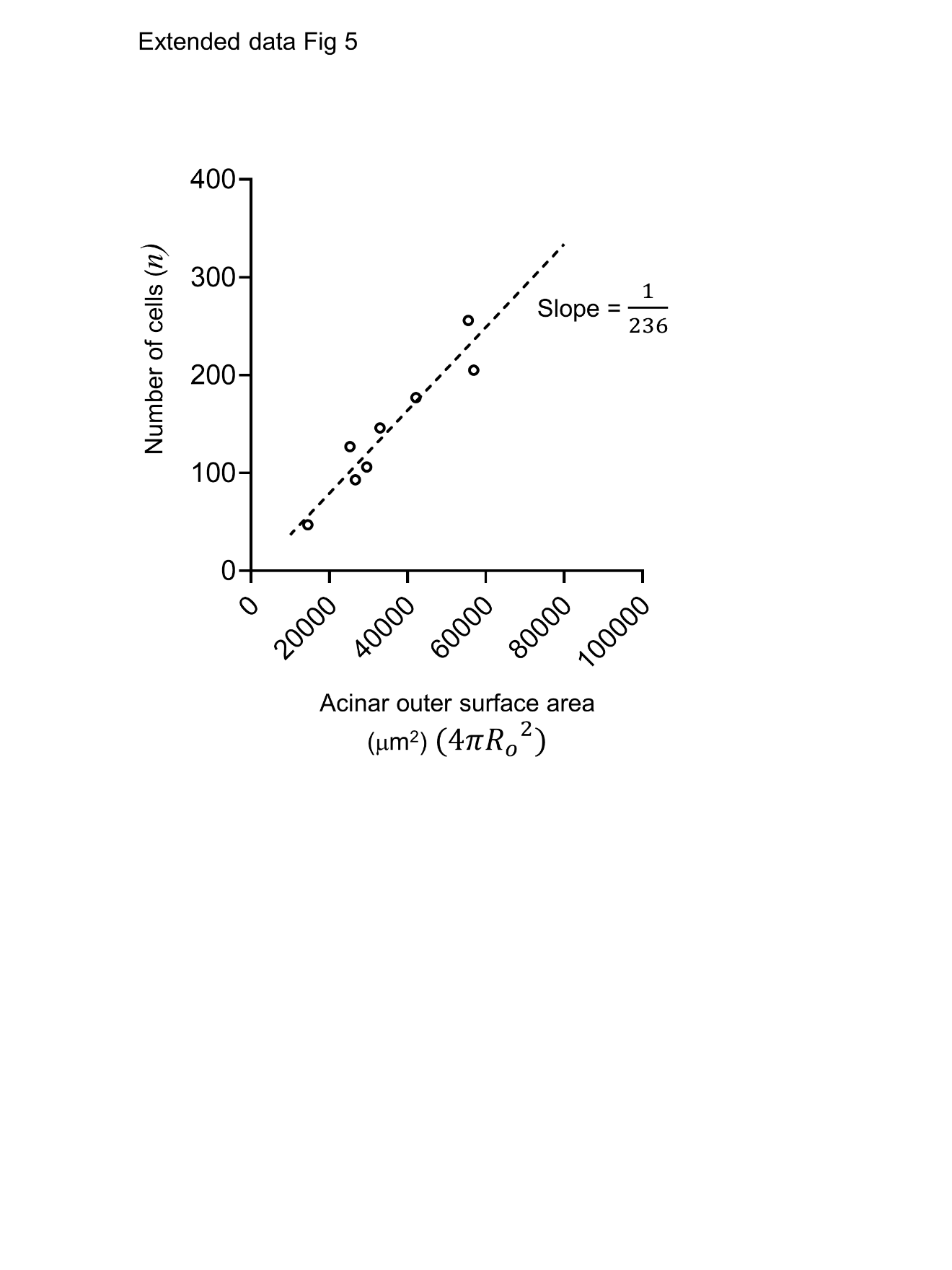

**Fig. S2.** Experimentally determined relationship between number of cells in acini (*n*) and acinar outer radii ($R_{o}$). Data shows *n* is directly proportional to ${R_{o}}^{2}$. The fitted line was obtained from measured values of *n* and outer surface areas ($4\pi{R_{o}}^{2}$) of 8 acini with a single lumen.

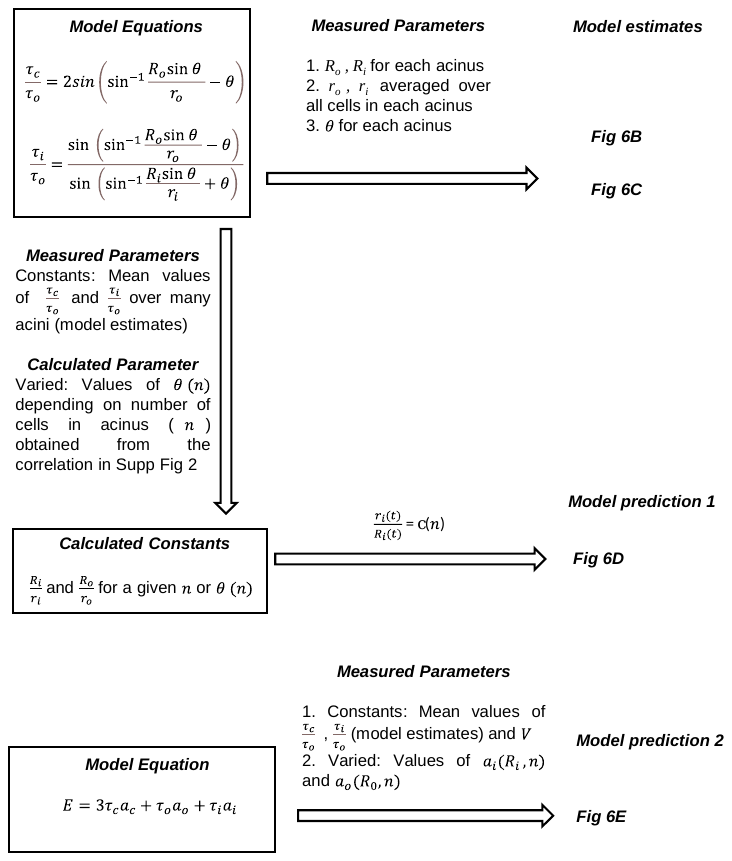

Fig S3. Schematic showing the equations and measured parameters used to obtain the model estimates and model predictions.

**Movie Legends**

**Movie 1.** Live imaging of MDCK acinus expressing GFP-podocalyxin formed over 10 days of three-dimensional culture. At time T = 0 h, the acinus was treated with Rho activator II. Scale bar, 20 μm.

**Movie 2.** Live imaging of MDCK acinus expressing GFP-podocalyxin formed over 10 days of three-dimensional culture and treated at time T = 0 h with Rho activator II. Scale bar, 20 μm.

**Movie 3.** Live imaging of MDCK acinus expressing GFP-podocalyxin formed over 10 days of three-dimensional culture and treated at time T = 0 h with Rho activator II. Scale bar, 20 μm.

**Movie 4.** Live imaging of MDCK acinus expressing GFP-podocalyxin after 10 days of three-dimensional culture and treated at time T = 0 h with DMSO as vehicle control. Scale bar, 20 μm.

**Movie 5.** Live imaging of MDCK acinus expressing GFP-H2B, emerald-occludin after 10 days of three-dimensional culture and treated at time T=0h with 1 μg/ml Rho Activator II. Scale bar, 25 μm.

**Movie 6.** 3D confocal scan of a single MDCK acinus expressing GFP-podocalyxin, treated with 1 μg/ml Rho Activator II for 12 h.

**Movie 7.** Live imaging of a single acinus expressing mCherry-KASH1 and GFP-podocalyxin. Cells were cultured for 10 days in matrigel, then treated at T = 0 h with doxycycline to induce the expression of mCherry-KASH1. Scale bars, 20 μm.

**Movie 8.** Live imaging of a single acinus expressing mCherry-KASH1 and GFP-podocalyxin. Cells were cultured for 10 days in matrigel, then treated at T = 0 h with doxycycline to induce the expression of mCherry-KASH1. Scale bar, 20 μm.

**Movie 9.** Live imaging of a single acinus expressing mCherry-KASH1 and GFP-podocalyxin. Cells were cultured for 10 days in matrigel, then treated at T = 0 h with doxycycline to induce the expression of mCherry-KASH1. Scale bar, 20 μm.

**Movie 10.** Live imaging of a single acinus expressing mCherry-KASHI1ΔPPPL and GFP-podocalyxin. Cells were cultured for 10 days in matrigel, then treated at T = 0 h with doxycycline to induce the expression of mCherry- KASH1ΔPPPL. Scale bar, 20 μm.

**Movie 11.** Live imaging of MDCK acinus expressing GFP-podocalyxin, after 7 days of three-dimensional culture and treated at time T=0 h with 8 μg/ml β-1 integrin function blocking antibody (AIIB2). Scale bar, 20 μm.

**Movie 12.** Live imaging of a single acinus assembled by 344SQ lung cancer cells treated with 1 μg/ml of Rho activator II at 0 h. Scale bar is 50 μm.

**Movie 13.** Live imaging of MDCK acinus expressing GFP-podocalyxin, after 7 days of three-dimensional culture where the acinus was laser ablated at T=0 h. Scale bar, 25 μm.

**Movie 14.** Live imaging of MDCK acinus expressing GFP-podocalyxin, after 7 days of three-dimensional culture where the acinus was laser ablated at T=0 h. Scale bar, 25 μm.
